# How flat can a horse be? Exploring 2D approximations of 3D crania in equids

**DOI:** 10.1101/772624

**Authors:** Cardini Andrea, Marika Chiappelli

## Abstract

Quantitative analyses of morphological variation using geometric morphometrics are often performed on 2D photos of 3D structures. It is generally assumed that the error due to the flattening of the third dimension is negligible. However, despite hundreds of 2D studies, few have actually tested this assumption and none has done it on large animals, such as those typically classified as megafauna. We explore this issue in living equids, focusing on ventral cranial variation at both micro- and macro-evolutionary levels. By comparing 2D and 3D data, we found that size is well approximated, whereas shape is more strongly impacted by 2D inaccuracies, as it is especially evident in intra-specific analyses. The 2D approximation improves when shape differences are larger, as in macroevolution, but even at this level precise inter-individual similarity relationships are altered. Despite this, main patterns of sex, species and allometric variation in 2D were the same as in 3D, thus suggesting that 2D may be a source of ‘noise’ that does not mask the main signal in the data. However, the problem is complex and any generalization premature. Morphometricians should therefore test the appropriateness of 2D using preliminary investigations in relation to the specific study questions in their own samples. We discuss whether this might be feasible using a reduced landmark configuration and smaller samples, which would save time and money. In an exploratory analysis, we found that in equids results seem robust to sampling, but become less precise and, with fewer landmarks, may slightly overestimate 2D inaccuracies.

## 1. Introduction

### 1.1. 2D to 3D approximation in geometric morphometrics: what is the problem?

The quantitative study of morphological variation has not lost its centrality in evolutionary biology and has in fact lived a renaissance thanks to the development of geometric morphometrics (GMM) (Rohlf and Marcus, 1993; Adams et al., 2004; Cardini and Loy, 2013). Landmark-based GMM using Procrustes methods employs sets of anatomically corresponding points to quantify and compare the size and shape of biological forms (Cardini, 2013). Landmark data can be collected in a variety of ways, among which digitizing anatomical points on photos is very common (Cardini, 2014). However, because most biological structures are highly three dimensional, and exceptions such as fly wings or plant leaves are relatively rare, 2D analyses of 3D anatomical features inevitably introduce an error by flattening the third dimension. This was stressed in the early days of applied GMM by Roth (1993) and has recently been brought again to the attention of the morphometric community by Álvarez and Perez (2013) and (Cardini, 2014). For brevity, as in Cardini (2014), we will call the problem of approximating a three-dimensional object with a flat picture as the “Two to Three Dimensional approximation”, henceforth abbreviated with TTD.

The most obvious solution to TTD is to avoid the issue in the first place by collecting 3D data using digitizers (e.g., http://www.3d-microscribe.com) or three-dimensional models obtained with scanners or photogrammetric reconstructions. However, 3D data acquisition may raise practical issues in terms of speed and costs. Especially for costs, 3D photogrammetry is in fact interesting and increasingly applied in GMM, especially on mammals of both large (Evin et al., 2016) and small size (Giacomini et al., 2019). This technique, pioneered in GMM by Fadda et al. (1997), allows to combine pictures of different views of an object to reconstruct a 3D model of its surface, thus, in a sense, ‘transforming’ a simple and relatively inexpensive digital camera in an accurate 3D scanner. 3D photogrammetry has made big technological progresses and, as the software for the construction of the 3D models is either free (Falkingham, 2012) or cheap (e.g., https://www.agisoft.com/), it offers an effective alternative to more expensive devices (high-resolution scanners, 3D digitizers etc.). Nevertheless, because of the large number of pictures necessary for an accurate reconstruction, plus the often long computational time necessary for building a 3D model, the technique requires much more time than taking one or a few pictures of, for instance, a cranium in ventral, lateral and dorsal view. There is an inevitable trade-off between *pros* and *cons* of different methods for data acquisition (Álvarez and Perez, 2013; Cardini, 2014; Navarro and Maga, 2016; Buser et al., 2018) and a morphometrician may have to decide whether the better accuracy of 3D landmarks is more important than having potentially larger samples using 2D photos. The answer to what is best is unlikely to be general and will depend on many factors, which will be often specific to each study. Thus, for a rigorous evidence-based decision, a preliminary analysis to compare 2D and 3D results may be necessary (Cardini, 2014).

Since Álvarez and Perez (2013) and Cardini (2014), more attention has been paid to TTD in GMM studies and several papers have focused on different aspects of the problem. For instance, Bakkes (2017) focused on a related issue, which is not specific of 2D images, but is likely to be more important in studies using flat images of 3D objects. This is the effect of the orientation on repeatability. Repositioning a specimen before collecting the data will add a source of error that is negligible only if the standardization of its position relative to the camera is extremely precise. Indeed, he found that, although repeatability of 2D data was high for the symmetric component of shape, variability in a specimen orientation introduced important differences in patterns of asymmetric variation in basis capituli of three species of ticks. That asymmetric variation, even when large, may be particularly affected by 2D analyses of pictures, was also suggested by a recent study (Hedrick et al., 2019). The authors employed slightly different landmark configurations to capture the shape of dog-fish vaginas both in 2D and 3D. Despite a good correlation between the two types of data, they discovered that some results were strongly impacted by TTD. As in (Bakkes, 2017), the patterns of asymmetries were profoundly altered in the 2D dataset. Also, and even more worrying, group structure (reproductive *vs* non-reproductive females in the sample) was evident in a PCA of 3D shape, but totally absent when the same analysis was done in 2D. This indicates that a dominant pattern of shape variation, evident in 3D, was totally lost (or hidden) in 2D.

A poor TTD may differentially affect different types of analyses and can even vary across taxa in a study. This was the case in a comparison of 2D and 3D data measuring the morphology of the head in Oligocottinae fishes (Buser et al., 2018). Although the main directions of shape variation suggested similar patterns, the authors found (p. 806) that “in taxa where shape variation in the z-axis is high, the 2D shape variables show sufficiently strong distortion to influence the outcome of the hypothesis tests regarding the relationship between mouth size and feeding ecology”. This meant that, despite working at a macro-evolutionary level, and thus presumably dealing with fairly large differences among taxa, only 3D data supported a well established ecomorphological relationship between mouth sizes and the proportion of preys in the diet. Also, 2D data, unlike 3D ones, showed a strong divergence of one lineage, that happened to be the one with “the greatest degree of z-axis body depth within Oligocottinae” (p. 806).

That large differences are no guarantee of a negligible TTD is also the conclusion of another paper (Santana et al., 2019). These authors explored macro-evolutionary patterns using sophisticated comparative models across a vast range of chiropteran species and compared results from ventral, lateral and frontal 2D views of the cranium with 3D data. However, instead of using photos, they obtained the 2D data by simply removing one of the three coordinates of the 3D landmarks, after a convenient alignment, in order to approximate what one would get on real 2D pictures. Despite the macro-evolutionary level of their work, they found important differences in patterns of shape variation not only across 2D views, but also between 2D and 3D data. Variability in results between views might be simply related to the different aspects of anatomy being captured by each of them, and thus to their potentially differential evolutionary patterns and rates. In contrast, incongruencies between 2D and 3D should not happen, if 2D is a faithful representation of 3D, and were therefore attributed by the authors to the loss of information in 2D. However, in this specific case and similar ones, an intriguing question is whether particularly sophisticated models, which make a large number of assumptions, often hard to test in real datasets, might be more sensitive to the effects of TTD.

Indeed, TTD seems to be strongly specific to the data and questions being investigated and, with relatively flat structures, 2D data could be adequate or even outperform high resolution 3D images in a cost-benefit assessment. This was suggested in a quantitative trait loci (QTL) analysis of the mouse hemi-mandible (Navarro and Maga, 2016). In this study, the morphometricians compared 2D landmarks with 3D ones, using the same anatomical points, but also compared the former with 3D data augmented by a densely sampled set of semilandmarks on the surface of the hemi-mandibles. Although 3D data were more powerful to detect associations with genetic sequences, 2D landmarks recovered 17 QTLs compared to 19 found with 3D landmarks. The addition of almost 580 3D semilandmarks only increased the total number of QTL with four more new loci. Thus, the authors stressed that 2D morphometrics has benefits (simplicity and speed) in large phenotyping studies, even if accuracy is reduced, and concluded that using 3D data can in fact increase power, but (p. 1160) on the other hand “the congruence of … results pleads for robustness of our knowledge on the genetic architecture of the mouse mandible, built over a few decades and initially based on 2D imaging techniques”.

### 1.2. TTD in megafauna: an example using equids

Current research on TTD clearly points towards a complex picture of the problem. It will require many more studies before researchers can try any generalization on when 2D is more or less appropriate. The brief overview we provided indicates that there are clearly several interesting avenues to explore in relation to TTD. Among these, an important but less obvious one is how TTD may impact analyses of really large anatomical structures. Indeed, none of the TTD studies we know has focused on the megafauna, an arbitrary and heterogeneous group of animals, whose body mass is larger than 45-50 Kg (Barnosky, 2008). This group includes some of the most charismatic living and recently extinct species, such as, among mammals, for instance, lions and sabre tooth tigers, elephants and mammoths, most marine mammals and many others. Research on anatomical and ecomorphological variation of modern and past representatives of the megafauna, whose partial but rapid extinction on land at the end of the Pleistocene still poses an unresolved dilemma (Koch and Barnosky, 2006), attracts great interest. Indeed, large mammals and birds, but also ‘reptiles’ such as crocodyles and dinosaurs, have been the subject of innumerable morphological studies, including many 2D GMM analyses (e.g., Pierce et al., 2008; Amaral et al., 2009; Brombin et al., 2009; Figueirido et al., 2009; Brusatte et al., 2012; Christiansen, 2012; Loza et al., 2015; Meloro et al., 2017; Page and Cooper, 2017; Angulo-Bedoya et al., 2019).

To start filling this gap in 2D GMM research, we chose to investigate TTD in the living equids, a group which has been extensively studied by morphometricians, especially in relation to the evolution of the lineage (e.g., (Radinsky, 1984; Heck et al., 2018)) and the domestication of the horse (e.g., (Eisenmann and Baylac, 2000; Bignon et al., 2005; Heck et al., 2018)). We focused on adult crania, that in horses, zebras and wild donkeys, weighing up to several hundreds kilos, can be more than half a meter in length and, therefore, is much bigger than anything analysed in previous TTD studies. More specifically, we analysed ventral cranial variation at both micro- and macro-evolutionary levels, using the same configuration of landmarks in 2D and 3D. For the micro-evolutionary analysis, where differences are typically small, we employed the largest available species sample, consisting of more than 100 plains zebras. For the macro-evolutionary study, with larger differences, we studied all seven living species of equids. Thus, to assess TTD, we compared results from the two sets of data using a variety of approaches, ranging from simple data visualizations and tests, performed in parallel for 2D and 3D data (1-3), to correlational methods (4) and analyses in a ‘common shape space’ (5-6):

1. Graphical summaries. The congruence of 2D and 3D size and shape variation was first inspected using respectively box-plots and scatterplots of the first principal components (PCs; including PCs of mean shapes at the macro-evolutionary level).
2. Group mean differences. The magnitude and significance of group differences in size and shape were tested, and 2D results were compared with 3D ones. In plain zebras, we tested sexual dimorphism, whereas in the macro-evolutionary analysis we analysed species differences. For shape, we also assessed group (sex or species) cross-validated classification accuracy.
3. Allometry. The significance of allometry, the relationship between size and shape, was tested and static (within species using adults) allometric trajectories were compared across groups. As in 2), groups were sexes for the micro-evolutionary level, and species for the macro-evolutionary study. At the macro-evolutionary level, we also estimated evolutionary allometry using species means. Results of each test were again compared between 2D and 3D analyses.
4. Correlations. The correlation of size and shape data were calculated at all the different levels (micro- and macro-evolutionary and using either all specimens or species means). If 2D is a faithful representation of 3D, correlations should be very high.
5. Within the same data space, following the approach proposed by Cardini (2014) to bring 2D and 3D data in the same shape space, we tested whether differences among individuals are larger than those between replicas using a hierarchical analysis of variance (ANOVA) (Klingenberg et al., 2002; Viscosi and Cardini, 2011; Cardini, 2014). Replicas here are 2D and 3D descriptions of size and shape of each specimen, and thus quantify the difference between the two types of data.
6. Finally, using shape data in the same Procrustes space as in 5), we built phenograms (all individuals, for plain zebras and all equids, and species mean shapes, for the macro-evolutionary level) to verify if 2D and 3D data of each specimen clustered as pairs in the trees, as it should happen if TTD is good.

In all analyses, as well as in the interpretation of results, as in previous studies by us or other researchers, the implicit assumption is that 3D is more accurate than 2D and, therefore, that smaller differences between the two types of data suggest a better TTD and thus a higher relative 2D accuracy.

## 2. Methods

### 2.1. Samples, data and geometric morphometrics

Data were collected in museums (see Acknowledgements for a list of institutions) using only adult animals with fully erupted molars. Specimens (Table 1) came both from the wild as well as from zoos. Zoo specimens were included to increase sampling in species poorly represented in museum collections. However, individuals with evident abnormalities were not digitized and potential outliers had been excluded in a previous study (Cardini, 2019). For the classification, we relied on museum catalogues, which, for plains zebras (*E. quagga* Boddaert, 1785), still reported the old classification as *E. burchelli*. Table 1 shows the sample composition, which is clearly heterogeneous and includes a fairly large sample only for plains zebras, our main dataset for the micro-evolutionary analyses. Seven specimens were of unknown sex, with canine size, often a clear dimorphic trait in equids, not providing unequivocal evidence to sex the animals. These few unsexed individuals were therefore excluded in all analyses of sex differences.

**Fig. 1.**
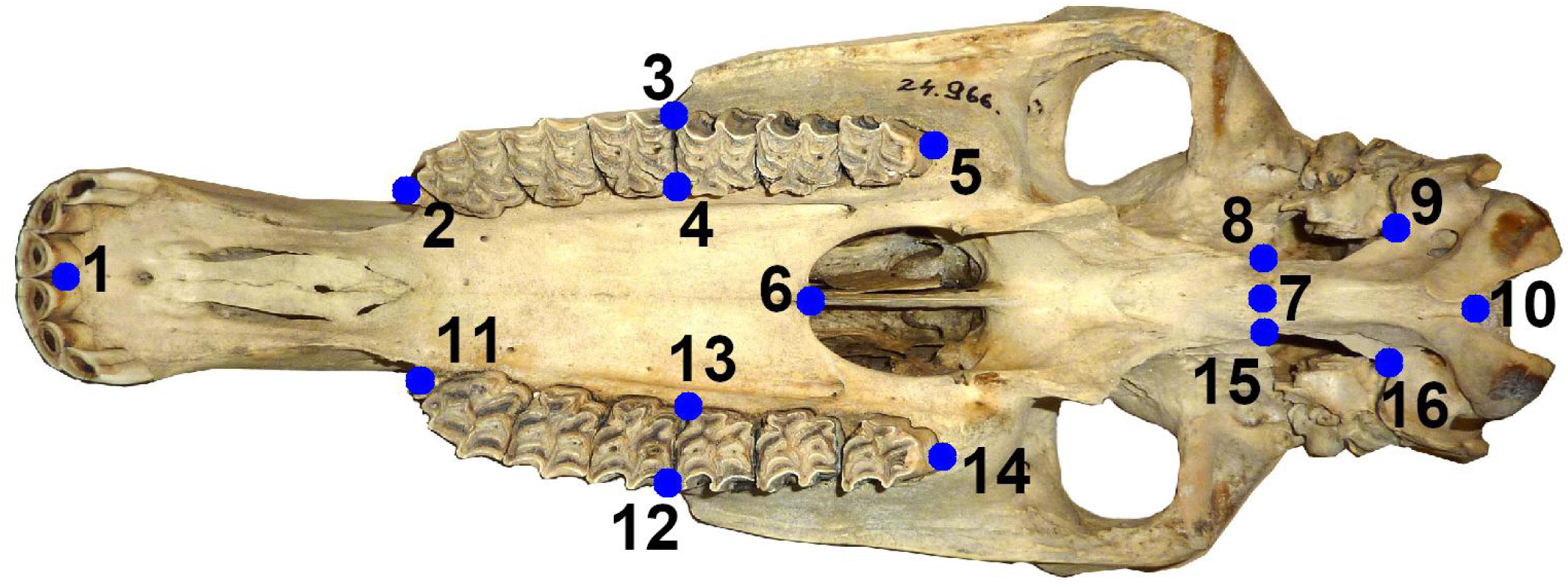
Landmark configuration on ventral crania of equids.

**Table 1.**
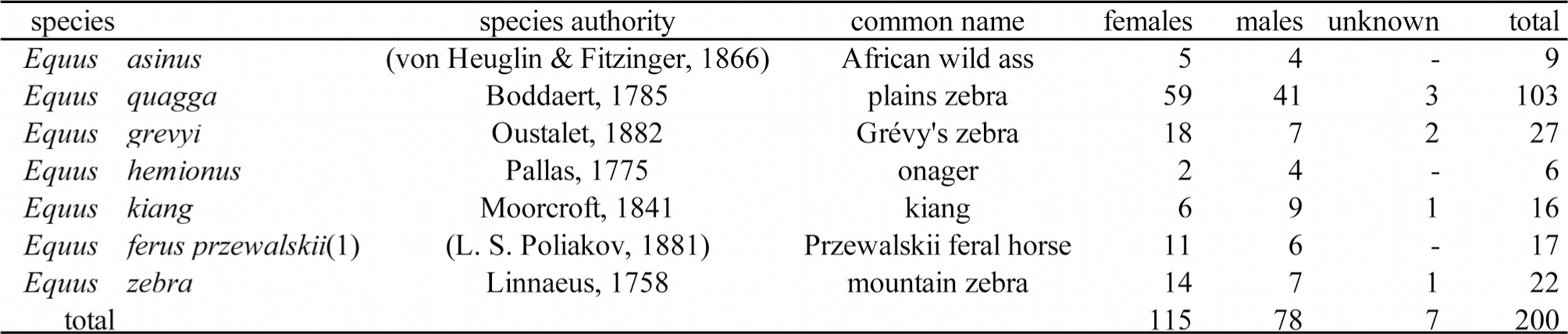
(1) A recent study suggests that living Przewalskii horse might in fact descend from an ancient group of domestic horses (Gaunitz et al., 2018). For brevity, in the figures we will refer to this sample simply as E. Przewalskii.

| species | species authority | common name | females | males | unknown | total |
| --- | --- | --- | --- | --- | --- | --- |
| *Equus asinus* | (von Heuglin & Fitzinger, 1866) | African wild ass | 5 | 4 | - | 9 |
| *Equus quagga* | Boddaert, 1785 | plains zebra | 59 | 41 | 3 | 103 |
| *Equus grevyi* | Oustalet, 1882 | Grévy's zebra | 18 | 7 | 2 | 27 |
| *Equus hemionus* | Pallas, 1775 | onager | 2 | 4 | - | 6 |
| *Equus kiang* | Moorcroft, 1841 | kiang | 6 | 9 | 1 | 16 |
| *Equus ferus przewalskii*(1) | (L. S. Poliakov, 1881) | Przewalskii feral horse | 11 | 6 | - | 17 |
| *Equus zebra* | Linnaeus, 1758 | mountain zebra | 14 | 7 | 1 | 22 |
| total |  |  | 115 | 78 | 7 | 200 |

3D Cartesian coordinates of anatomical landmarks were collected using a Microscribe 3D digitizer (http://www.3d-microscribe.com/). Because the 3D data were obtained from a previous study with a different focus (Cardini, 2019), 3D landmarks were available only on the left side of the cranium and were mirror reflected to reconstruct the missing right side (Cardini, 2016). Analyses on a range of mammals have shown that the inaccuracy introduced by using crania measured only on one side (an expedient to speed up data collection and increase sample size) is negligible (Cardini, 2017), if one is not interested in studying asymmetries and a structure has no evident directional differences between sides (Cardini, 2016).

From the original configuration of 3D landmarks on the entire cranium, we selected a subset of points that are clearly visible in pictures of ventral crania and are roughly coplanar (Fig. 1). On these pictures, landmarks were digitized in TPSDig (Rohlf, 2015). The photos were taken with a Panasonic Lumix DMC TZ6 camera that was held approximately parallel to the ventral surface of each cranium positioned upside down on a table. The distance and zoom of the camera varied and there might be small photographic distortions, which are therefore incorporated in the 2D error. As in previous TTD studies, differences between 2D and 3D include, in fact, also the precision in landmark digitization. However, it is likely that both small photographic distortions and landmarking errora are of modest size compared to the variation introduced by the 2D flattening of a 3D structure. For instance, Cardini (2014) showed that the error due to 2D flattening was almost 4 times larger than the sum of differences related to positioning, digitizing and photographic device, and he found this in marmot hemi-mandibles, which are flat and, in terms of TTD, perform much better than crania. Nevertheless, although we do not mention these sources of error further, readers should bear in mind that they are part of our total estimate of measurement error and they may, therefore, slightly inflate the magnitude of TTD. Also, because the scale factor included in the pictures was not always consistently positioned on the ventral surface of the cranium in all specimens, we used the 3D measurement of condylobasal length to rescale the 2D landmarks.

From the original Cartesian coordinates of the landmarks, size was measured using centroid size (CS - i.e., the square root of the sum of the distances of each landmark from its centroid) and shape was computed using a Procrustes superimposition to remove size variation and minimize translational and rotational differences (Adams et al., 2004). On pictures we landmarked both sides and later discarded small asymmetries in shape (Klingenberg et al., 2002) to compare 2D and 3D data. Estimates of 2D size and shape based on the full configuration or just, as in 3D, its left half were almost perfectly correlated (size r=1.00, shape r=0.97). Thus, the small inconsistency in the protocol of 2D and 3D data collection (one using both sides and the other only the left half) has no impact at all on the outcome of the TTD analyses.

### 2.2. Statistical analyses

Size (CS) and Procrustes shape coordinates were computed in MorphoJ (Klingenberg, 2011) and imported in R (R Core Team, 2018), where we conducted most of the analyses. Details on methods are provided below using the same subdivision as in the Introduction. As anticipated, analyses 1-4) were performed using separate 2D and 3D Procrustes shape spaces, whereas analyses 5-6) were done in a common shape space using the method developed by Cardini (2014).

1) Box plots of CS were drawn in R (R Core Team, 2018) using ggplot (Wickham and Wickham, 2007) and compared between 2D and 3D data. The same R package was used for drawing scatterplots of the first shape PCs, which were computed in R using the prcomp function (R Core Team, 2018).

2) Mean group differences (sex, for plains zebra, and species, regardless of sex, for the macro-evolutionary analyses) were tested with ANOVAs using the adonis function of the vegan package (Oksanen et al., 2013)). Significance of size and shape was estimated using 10000 permutations of Euclidean distances and effect size was estimated using R^2^ (univariate for CS and multivariate for shape). R^2^ is the percentage of variance accounted for by the effect being tested. The adonis function was used also in all other tests of groups, as well as in the allometric analyses (see below).

The accuracy of shape for predicting groups (i.e., the classification according to sex or species, respectively, at the micro- and macro-evolutionary levels) was estimated using leave-out cross-validations and two types of functions. One employed Euclidean shape distances to classify individuals in groups using a between-group principal component analysis (bgPCA (Cardini et al., 2019), and references therein) in Morpho (Schlager, 2017). The other classification was obtained using Mahalanobis distances and a conventional leave-out linear discriminant function (DA) in MASS (Venables and Ripley, 2002)). Overall predictive accuracy was summarized in both analyses using the average cross-validated percentage of correctly classified individuals.

3) Allometric trajectories were calculated with multivariate regressions of shape onto CS (Klingenberg, 2016). Static allometries of adults were compared first between sexes in plains zebras and then across all equid species (regardless of sex) using a permutational MANCOVA (multivariate analysis of covariance, CS by group, with CS as covariate (Anderson M.J., 2001; Zelditch et al., 2004; Oksanen et al., 2013)). In this analysis, the interaction between size and group tests the significance of the divergence of the allometric trajectories. Finally, evolutionary allometry was estimated by regressing species mean shapes onto the corresponding mean CS. Regressions and MANCOVAs were done in vegan using the adonis function (Oksanen et al., 2013).

4) The congruence between 2D and 3D data was assessed also by computing correlations. For CS, we used the correlation coefficient and, for shape, the matrix correlation between the Euclidean shape distances computed pairwise between observations in each shape space (i.e., 2D and 3D). Matrix correlations were calculated with the mantel function in vegan (Oksanen et al., 2013).

5) To perform analyses in the same shape space, which allow to use the standard GMM ANOVA protocol (see below) for the assessment of measurement error (Arnqvist and Martensson, 1998; Klingenberg et al., 2002; Viscosi and Cardini, 2011; Fruciano, 2016), we followed the method proposed by Cardini (2014): thus, we added a fake Z coordinate (equal to zero) to the 2D landmarks; merged the data with the 3D ones, and did a common 3D superimposition; and, in MorphoJ (Klingenberg, 2011), regressed the resulting shape coordinates onto a dummy variable coding for the type of data (i.e., 2D *vs* 3D) to compute residuals. Regression residuals are equivalent to mean centered 2D and 3D data, an operation that could be done manually by subtraction (2D shapes minus their mean and 3D shapes minus the 3D mean). The advantage of using the residuals from MorphoJ is that this software automatically adds them to the grand mean of the data, so that the resulting coordinates can be reimported and used as shape data in this or other programs. Mean centering is an expedient to control for the bias due to the lack of a real Z coordinate (measuring depth) in the 2D data, but it does not change the relative shape distances within each dataset, because the quantity removed is in both cases a constant. This is easily verified by computing the matrix correlations of the 2D residuals with the original 2D shapes and by doing the same for the 3D data (both producing r=1.000). Cardini (2014) provides more details on the ‘mean centered-common shape space approach’, as well as example data to replicate the analysis.

In plains zebras (micro-evolutionary level), the mean centered data in the common Procrustes shape space were analysed using a hierarchical multivariate ANOVA (MANOVA) with sex as main factor and individual as random factor (Arnqvist and Martensson, 1998; Fruciano, 2016). Using the same design and type of ANOVA, but adding first species as a main factor (followed by sex and individuals), the analysis was repeated at the macro-evolutionary level including all species and specimens of known sex. The ANOVAs, performed also for CS, allowed to re-test in a common data space the main effects of species and sex, but also, and more importantly, to assess whether differences among individuals were significantly larger than those between replicas (2D *vs* 3D), our estimates of measurement error in relation to (mainly) 2D inaccuracies.

6) Finally, UPGMA (unweighted pair group method with arithmetic mean) phenograms were computed by applying the hclust function in R (R Core Team, 2018) to the matrix of Euclidean shape distances computed pairwise in the common shape space. For the micro-evolutionary analysis, the phenogram was computed including only the 103 specimens of plains zebras. For the macro-evolutionary study, phenograms were computed both using all 200 specimens as well as the species mean shapes.

## 3. Results

Results are presented following the same order and numbering as in the Introduction and method sections on statistical analyses.

### 3.1. Graphical summaries

Figure 2 summarizes the patterns of variation in size within plains zebras, using separate sexes, and across species, with pooled sexes. CS is slightly (ca. 2% or less on average) underestimated in 2D data, but the congruence with 3D data is otherwise striking. For shape (Figs 3-4), the comparison of scatterplots of 2D and 3D data is less straightforward, but suggests fairly good congruence especially at macro-evolutionary levels (Fig. 4). Within plains zebras, 2D and 3D scatterplots of shape PCs indicate both a complete overlap of females and males (Fig. 3), although in 2D data PC1 is more stretched and females appear to vary more than males. In the interspecific analysis, species overlap partially (Fig. 4), with confidence ellipses suggesting a very similar pattern of relative differences in 2D and 3D. As it had occurred at the micro-evolutionary level, also including all specimens and species, PC1 accounts for more variance in 2D than in 3D.

**Fig. 2.**
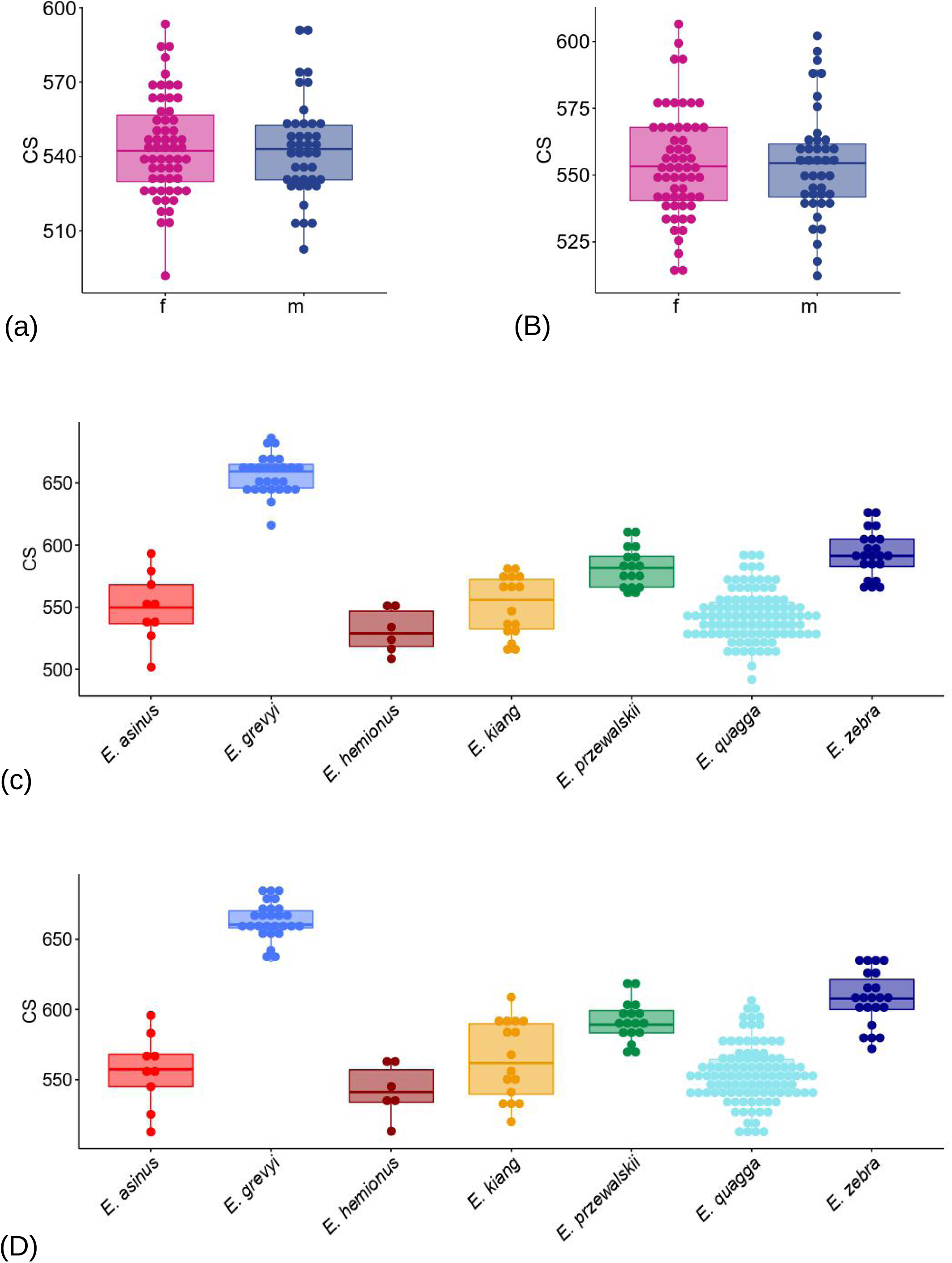
Box and jitter plots of CS in females (left, pink colour) and males (right, blue colour) of plains zebras (A, 2D and B, 3D) and in the total sample subdivided by species regardless of sex (C, 2D and D, 3D).

**Fig. 3.**
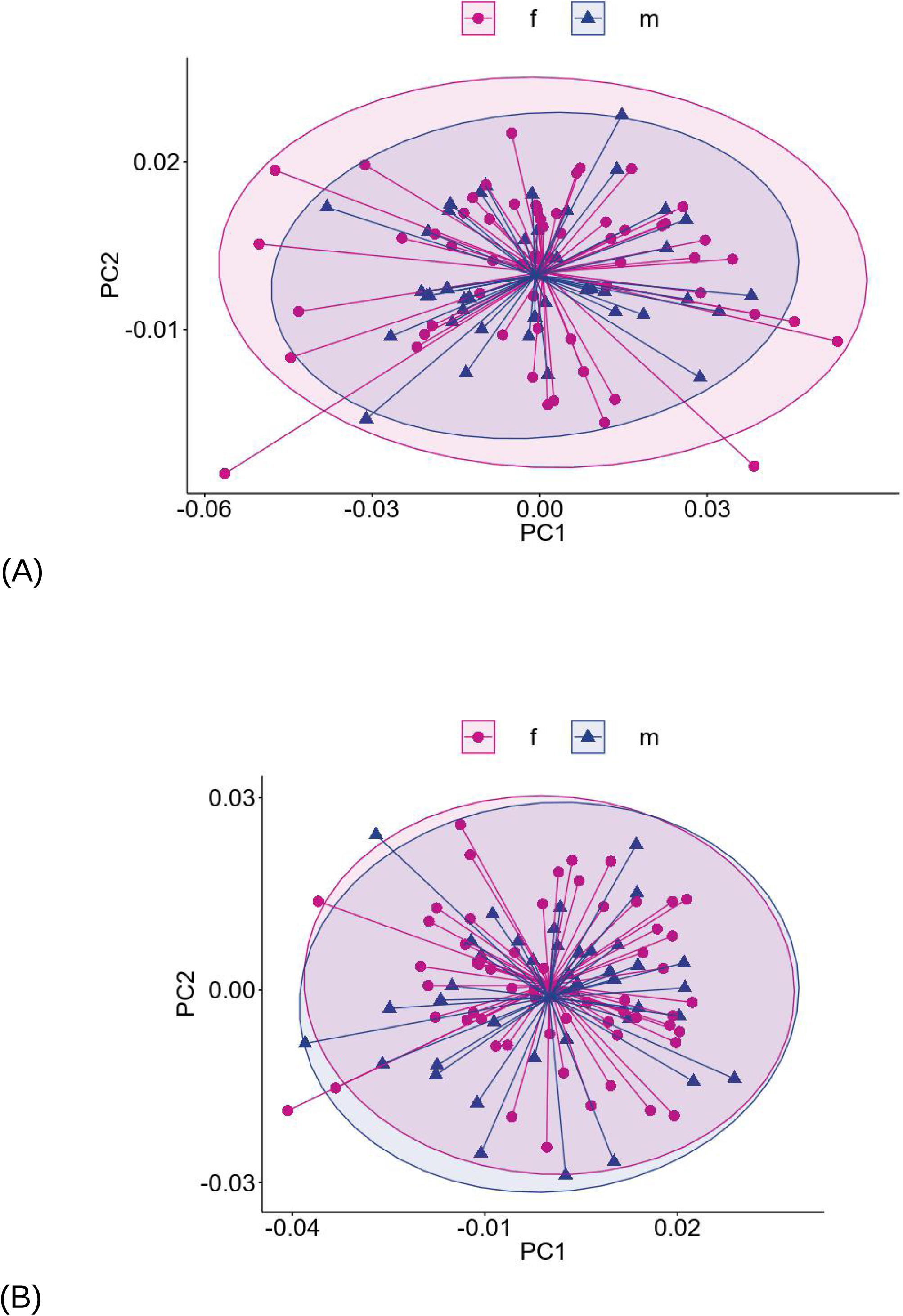
Separate shape spaces: scatterplots of PC1-PC2 with 95% confidence ellipses for females (pink) and males (blue) in 2D (A, variance accounted for by PC1-2: 21-13%) and 3D (B, variance accounted for by PC1-2: 15-12%).

**Fig. 4.**
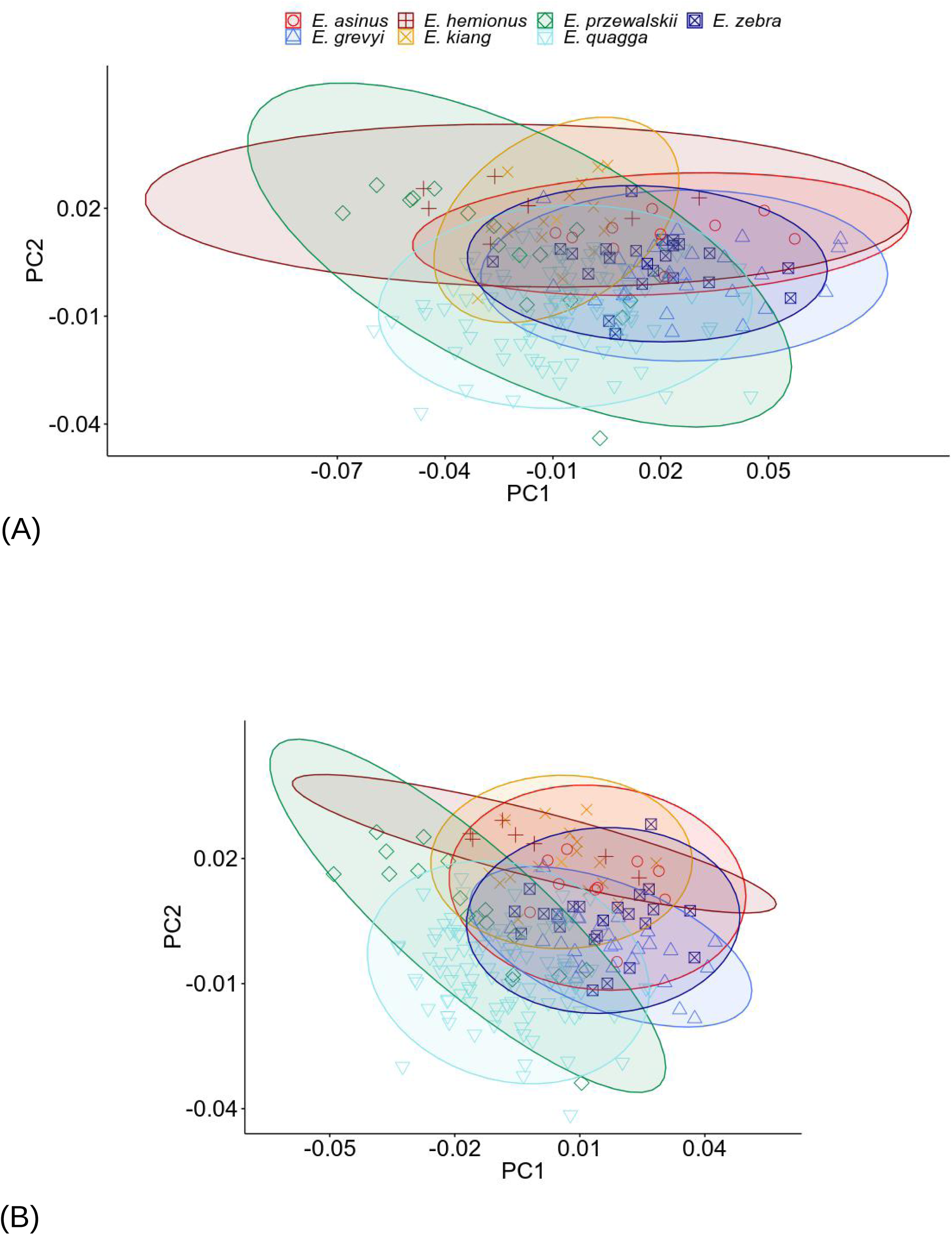
Separate shape spaces: scatterplots of PC1-PC2 with 95% confidence ellipses for different species (regardless of sex) in 2D (A, variance accounted for by PC1-2: 26-14%) and 3D (B, variance accounted for by PC1-2: 18-14%).

### 3.2. Group mean differences

Results of the tests of group mean differences run in parallel with data in separate spaces are shown in Table 2. In plains zebras (Table 2A), sexual dimorphism is totally absent in size (R^2^<0.1%) and very small in shape (R^2^<2%). This result is the same in 2D and 3D, and congruent with the patterns suggested by the box-plots (Fig. 2a-b) and PCAs (Fig. 3). Sex classification accuracy based on shape is virtually identical in 2D and 3D and negligibly better (ca. 60%) than an approximate 50% random chance expected for two groups of almost equal size. Thus, bgPCA and DA both confirm the absence of significant sexual dimorphism in ventral crania.

**Table 2.**
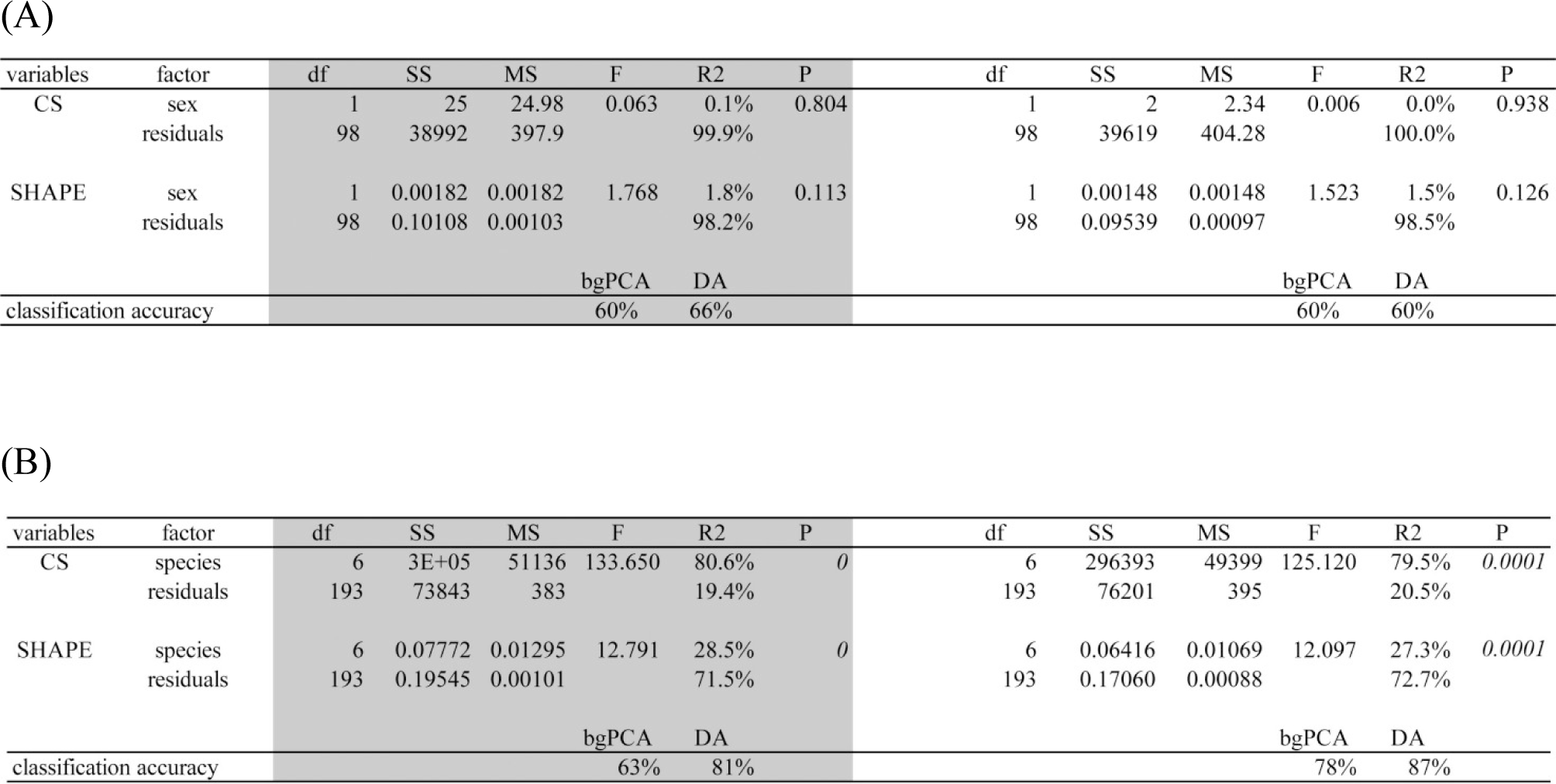
Group (sex or species) differences tested using permutational ANOVAs for size and shape and cross-validated classification accuracy estimated using bgPCAs and DAs of shape data to predict group affiliation. (A) Micro-evolutionary level testing sexual dimorphism in plains zebras. (B) Macro-evolutionary level testing inter-specific differences. (Here and elsewhere in the tables P<0.05 are in italics and, in analyses using separate data spaces, 2D results are emphasized with a grey background).

Also at the macro-evolutionary level (Table 2B), results of tests for species differences in 2D and 3D are almost identical for both size and shape, with highly significant differences accounting for 80% of variance in size and slightly less than 30% in shape. Species average classification accuracy using shape is about 80% (regardless of method and type of data) with the exception of the bgPCA, that produces a lower accuracy (63%), but still one which is much higher than expected by random chance with seven groups (∼14%, if they had the same size). Thus, despite overlaps in the PCAs (Fig. 4), equids show significant ventral cranial differences and, consistently with similarities in both box-plots of size and scatterplots of shape, the pattern is almost identical in 2D and 3D data.

### 3.3. Allometry

As for group differences at both evolutionary levels, analyses of allometry produce similar results in 2D and 3D (Table 3). In plains zebras (Table 3A), static allometries are significant in both females and males, and account for approximately 5-10% of variance (slightly more in females than males), whereas slopes do not differ significantly between sexes. R^2^ of sex is small and intercepts are non-significant or marginally significant after removing the interaction in the MANCOVA (results not shown). Thus, both 2D and 3D data lead to the same conclusion: a small amount of static allometric variation in plains zebras, with almost overlapping trajectories in females and males.

**Table 3.** Tests of static and evolutionary allometry. (A) Static allometry at micro-evolutionary level: sex by CS MANCOVA of shape, and multivariate regressions of shape on CS using separate sexes. (B) Macro-evolutionary level: species by CS MANCOVA of shape (static allometry), and multivariate regression of mean species shape onto mean species CS (evolutionary allometry).

| group | factor | df | SS | MS | F | R2 | P | df | SS | MS | F | R2 | P |
| --- | --- | --- | --- | --- | --- | --- | --- | --- | --- | --- | --- | --- | --- |
| both sexes | CS | 1 | 0.00805 | 0.00805 | 8.384 | 7.8% | 0 | 1 | 0.00516 | 0.00516 | 5.508 | 5.3% | 0.0001 |
|  | sex | 1 | 0.00187 | 0.00187 | 1.944 | 1.8% | 0.093 | 1 | 0.00148 | 0.00148 | 1.578 | 1.5% | 0.111 |
|  | sex by CS | 1 | 0.00081 | 0.00081 | 0.843 | 0.8% | 0.471 | 1 | 0.00039 | 0.00039 | 0.415 | 0.4% | 0.96 |
|  | meas. error | 96 | 0.09218 | 0.00096 |  | 89.6% |  | 96 | 0.08985 | 0.00094 |  | 92.8% |  |
| female | CS | 1 | 0.00720 | 0.00720 | 7.052 | 11.0% | 0 | 1 | 0.00393 | 0.00393 | 4.080 | 6.7% | 0.0002 |
|  | meas. error | 57 | 0.05816 | 0.00102 |  | 89.0% |  | 57 | 0.05487 | 0.00096 |  | 93.3% |  |
| male | CS | 1 | 0.00171 | 0.00171 | 1.957 | 4.8% | 0.081 | 1 | 0.00161 | 0.00161 | 1.795 | 4.4% | 0.067 |
|  | meas. error | 39 | 0.03402 | 0.00087 |  | 95.2% |  | 39 | 0.03498 | 0.00090 |  | 95.6% |  |

(B)
| group | factor | df | SS | MS | F | R2 | P | df | SS | MS | F | R2 | P |
| --- | --- | --- | --- | --- | --- | --- | --- | --- | --- | --- | --- | --- | --- |
| species | CS | 1 | 0.04006 | 0.04006 | 42.164 | 14.7% | 0 | 1 | 0.02129 | 0.02129 | 25.055 | 9.1% | 0.0001 |
|  | species | 6 | 0.04749 | 0.00792 | 8.332 | 17.4% | 0 | 6 | 0.04795 | 0.00799 | 9.404 | 20.4% | 0.0001 |
|  | species by CS | 6 | 0.00891 | 0.00148 | 1.563 | 3.3% | 0.055 | 6 | 0.00743 | 0.00124 | 1.458 | 3.2% | 0.0300 |
|  | meas. error | 186 | 0.17671 | 0.00095 |  | 64.7% |  | 186 | 0.15808 | 0.00085 |  | 67.3% |  |
| mean species | CS | 1 | 0.00142 | 0.00142 | 2.837 | 36.2% | 0.06 | 1 | 0.00077 | 0.00077 | 1.707 | 25.5% | 0.136 |
|  | meas. error | 5 | 0.00250 | 0.00050 |  | 63.8% |  | 5 | 0.00227 | 0.00045 |  | 74.5% |  |

The results of the MANCOVAs for differences of species static allometries are also highly congruent between 2D and 3D data. Differences in slopes are small but significant and the effect of species is significant and large, which overall suggests clearly distinct and modestly divergent trajectories. Evolutionary allometry, tested using species means, does not reach significance, but is large both in 2D and 3D (respectively, with R^2^ of 36% and 26%) and the pattern of allometric shape variation is very similar (Fig. 5).

**Fig. 5.**
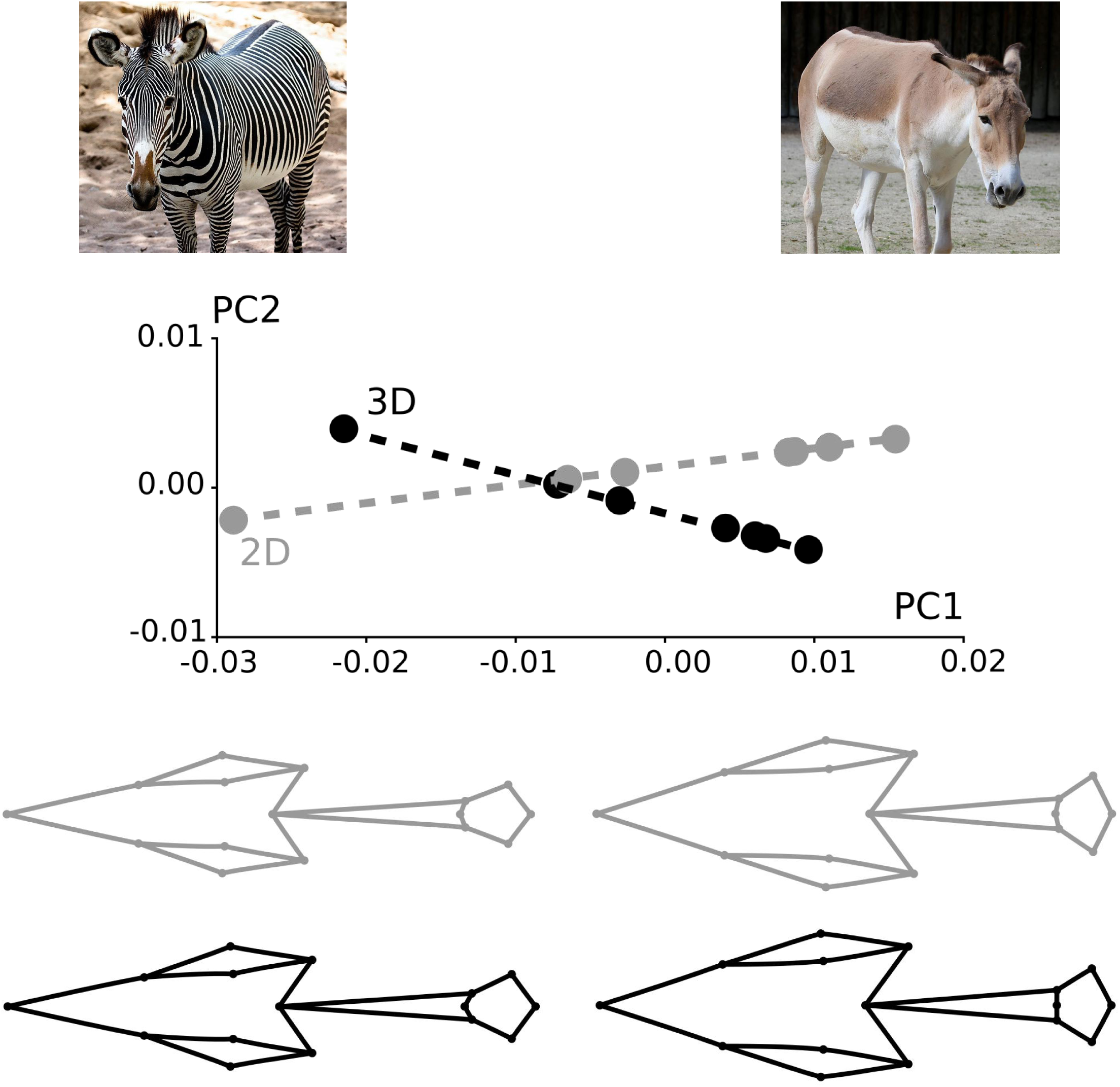
Evolutionary allometric variation in 2D (grey) and 3D (black), with wireframe visualization (Klingenberg, 2013) of ventral views predicted by the allometric trajectories at the opposite extremes of species mean size variation (magnified twice to aid comparisons). Opposite extremes correspond to the smallest and largest species, which are respectively the onager and Grevy’s zebra (pictures modified from https://en.wikipedia.org/wiki/File:Onager_Asiatischer_Wildesel_Equus_hemionus_onager_Zoo_Augsburg-10.jpg - licensed under the Creative Commons Attribution-Share Alike 3.0 Unported license – and from https://commons.wikimedia.org/wiki/File:Grevy%E2%80%99s_Zebra.jpg - available under the Creative Commons CC0 1.0 Universal Public Domain Dedication). Although allometric analyses were done in separate shape spaces, as a more effective and concise summary of the results, we visualized here the trajectories using a PCA of the predicted allometric shapes (Adams and Nistri, 2010) in the common space of 2D and 3D data (variance accounted for by PC1-2: 92.0-4.3%).

**Fig. 6.**
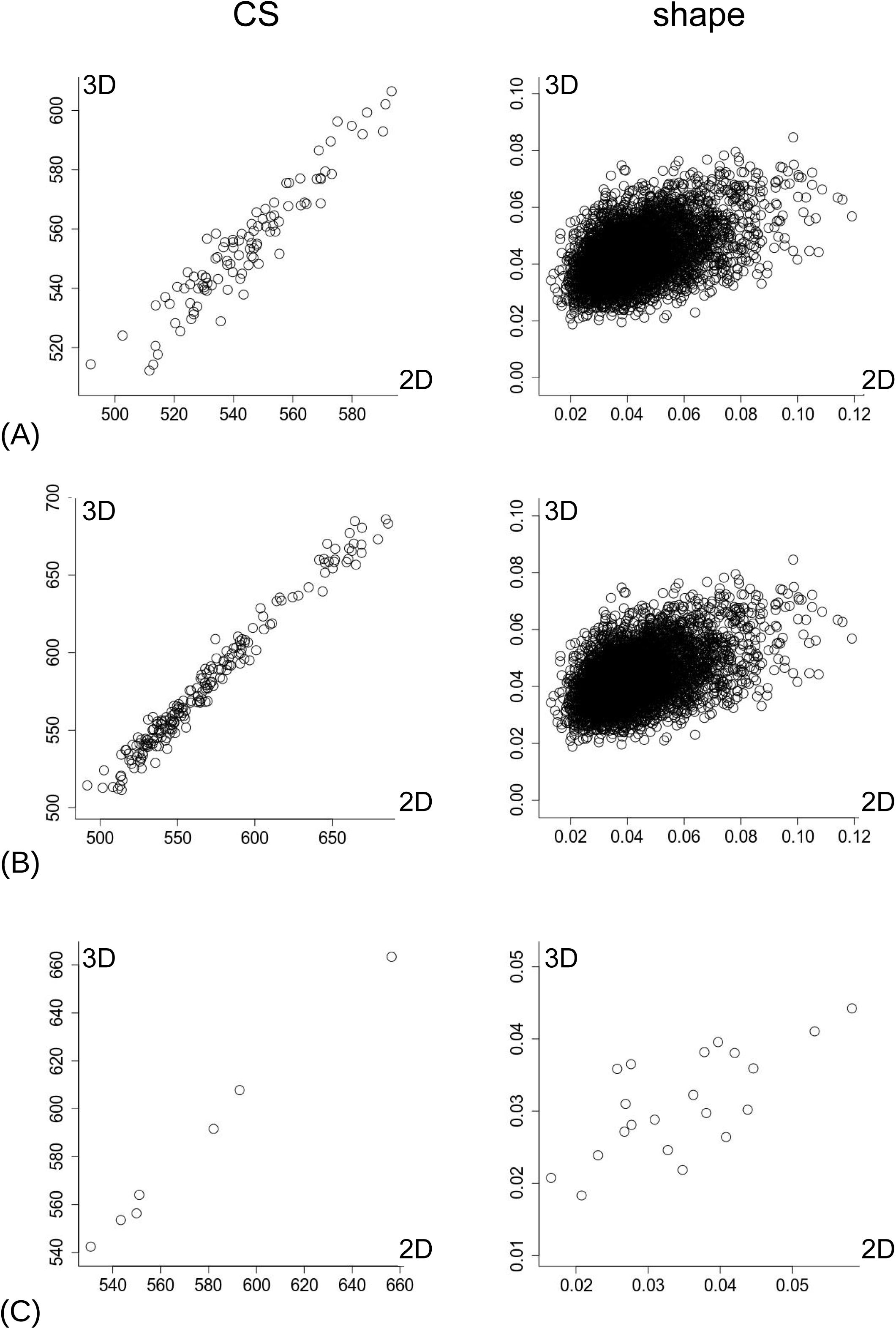
Separate shape spaces: scatterplots of CS (left column) and pairwise shape distance matrices (right column) from 2D (horizontal axis) and 3D (vertical axis) data: A) plains zebras; B) all individuals and species; C) species means.

### 3.4. Correlations

Correlations of 2D with 3D estimates of size and shape are shown in Table 4. As expected from previous analyses and especially from the strong similarities in the box-plots (Fig. 2), the correlations between estimates of CS in 2D and 3D are very high at all levels, ranging from 0.95 to almost 1 at respectively micro- and macro-evolutionary levels. In contrast, correlations of shape distances in the 2D and 3D spaces are much smaller, with plains zebras slightly below 0.5 whereas correlations at macro-evolutionary level vary between ca. 0.6 (all specimens and species) and 0.7 (species means).

**Table 4.** Correlations of size and shape (distances) at micro- and macro-evolutionary levels.

| level | N | CS | SHAPE |
| --- | --- | --- | --- |
| plains zebras | 103 | 0.95 | 0.47 |
| all species | 200 | 0.99 | 0.61 |
| species means | 7 | 1.00 | 0.71 |

### 3.5. ANOVAs in the common shape space

Table 5 shows the results of the ANOVAs in the common shape space, after mean centering the two datasets. Because micro- (plains zebras) and macro-evolutionary analyses produced similar results, we mainly focus on the more inclusive macro-evolutionary level (Table 5B). For size, the effect of species is very large (almost 80% of variance explained), sex is totally negligible and individuals explain about 10 times more variance than measurement error (ca. 90% vs 9% and ca. 20% vs 2% of variance in respectively plains zebras and all species). For shape, the effect of species is highly significant and accounts for slightly more than 25% of variance. The effect of sex is, in contrast, very small (R^2^ ca. 1-1.5%), but statistically significant. This seems surprising, as the size of the effect (R^2^) is very small and previous tests in separate spaces (2) were never significant. As it is a minor point, we clarify here the reason for this, likely spurious, incongruence. The F test statistics in the ANOVA has the advantage of simplicity and is generally robust, but it assumes isotropic variation around landmarks, which is a poor approximation for biological data (Klingenberg et al., 2002). Other test statistics, such as Pillai’s trace, do not make this assumption and are potentially more accurate, but they are not available in the permutational ANOVA we used. However, the F statistics (parametric or based on permutations) and Pillai’s trace (available as a parametric test in MorphoJ) generally produce congruent P values. In this specific case, in contrast, sex is significant using F but not using Pillai’s trace (not shown); as Pillai’s trace does not assume isotropy and is in agreement with non-signficant results of the permutation tests in the separate shape spaces (2) (and also consistent with a tiny R^2^ for sex), it seems reasonable that the F test for sex was unreliable.

**Table 5.** ANOVA in common data space at micro- (A) and macro-evolutionary (B) levels.

| level | variables | factor | df | SS | MS | F | R2 | P |
| --- | --- | --- | --- | --- | --- | --- | --- | --- |
| (A) | CS | sex | 1 | 6 | 6.02 | 0.080 | 0.0% | 0.7734 |
|  |  | individuals | 98 | 76510 | 780.7 | 10.326 | 91.0% | 0.0001 |
|  |  | residuals | 100 | 7560 | 75.6 |  | 9.0% |  |
|  | SHAPE | sex | 1 | 0.00302 | 0.00302 | 6.295 | 1.5% | 0.0002 |
|  |  | individuals | 98 | 0.15035 | 0.00153 | 3.201 | 74.7% | 0.0001 |
|  |  | residuals | 100 | 0.04793 | 0.00048 |  | 23.8% |  |
| (B) | CS | sp | 6 | 563837 | 93973 | 1202.850 | 78.2% | 0.0001 |
|  |  | sex | 1 | 170 | 170 | 2.180 | 0.0% | 0.1450 |
|  |  | individuals | 185 | 141725 | 766 | 9.810 | 19.7% | 0.0001 |
|  |  | residuals | 193 | 15078 | 78 |  | 2.1% |  |
|  | SHAPE | sp | 6 | 0.12506 | 0.02084 | 44.305 | 25.5% | 0.0001 |
|  |  | sex | 1 | 0.00422 | 0.00422 | 8.969 | 0.9% | 0.0001 |
|  |  | individuals | 185 | 0.26948 | 0.00146 | 3.096 | 55.0% | 0.0001 |
|  |  | residuals | 193 | 0.09080 | 0.00047 |  | 18.5% |  |

The most interesting effect tested in the ANOVA is, however, individual shape variation compared to differences between 2D and 3D shapes. This factor is highly significant and accounts for about three times more variance than the differences between the two types of data. Despite this, TTD errors are clearly large for shape, as they account for about 1/5 (all species included) to ¼ (plains zebras) of sample variance.

### 3.6. Phenograms in the common shape space

The effect of the large magnitude of TTD errors in shape becomes evident in the phenograms using individuals. The trees are better able to capture small differences between specimens compared to a PCA, but, with ca. 200-400 observations in the common 2D-3D space, they cannot be easily shown in a figure. The pattern they suggest is, however, very clear and can be summarized in words: only nine out of 103 specimens of plains zebras (9%) and only 16 out of 200 individuals (8%) in the total equid sample have 2D and 3D shape replicas of an individual clustering as ‘sisters’ in the phenograms. This indicates a rather poor correspondence, if one inspects the precise inter-individual similarity relationships of 2D and 3D shapes. Only with mean shapes (Fig. 7), 2D and 3D data of each species always cluster together as ‘sisters’, which supports the observation from the correlational analyses (4) that species means show the highest congruence between datasets and thus the best TTD.

**Fig. 7.**
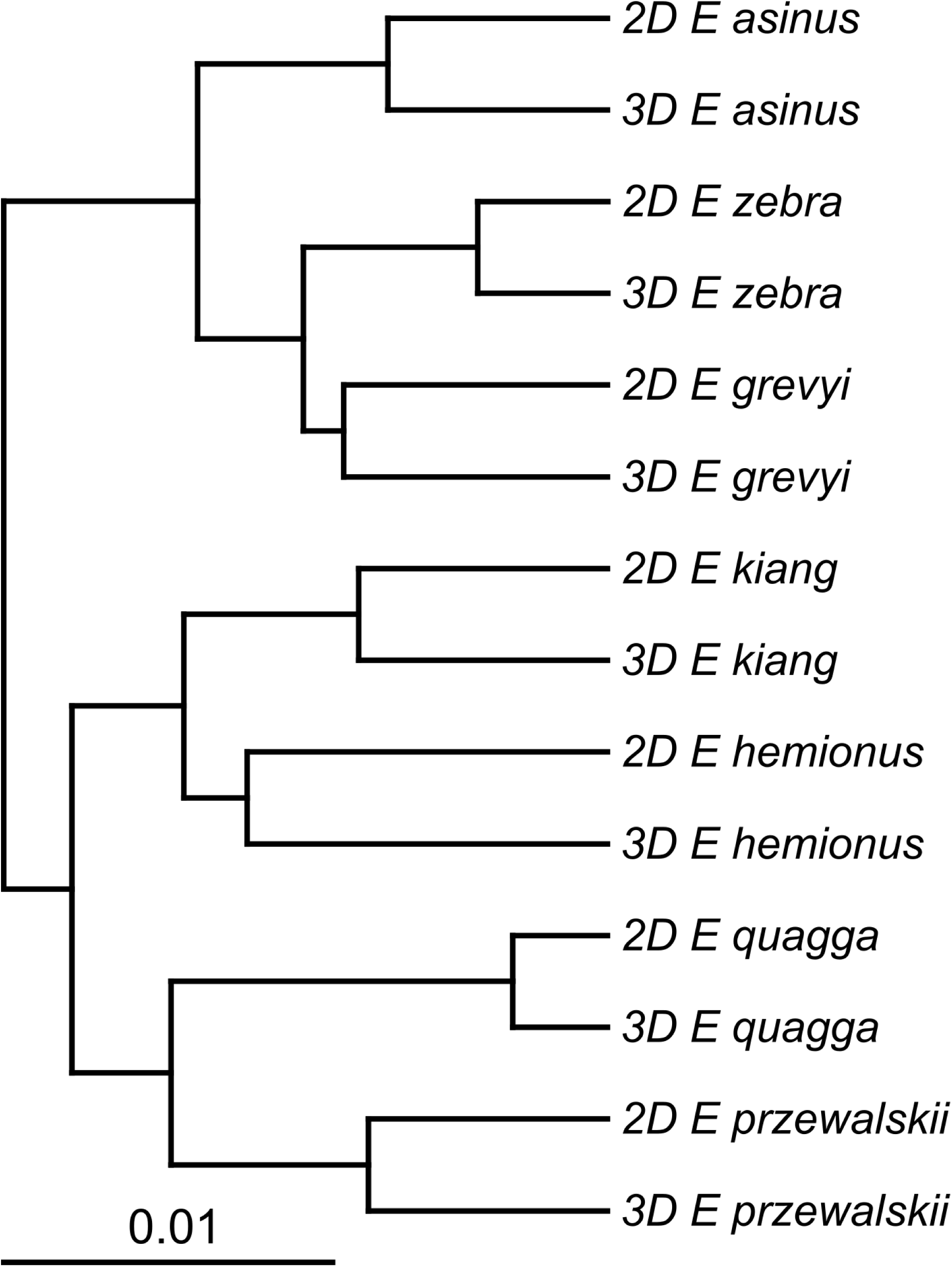
Common shape space mean-centered 2D-3D data analysis: UPGMA phenogram of species mean shapes.

## 4. Discussion

Before moving to the main conclusions from our TTD study, we will touch on a number of smaller, but nevertheless important points and methodological issues. As size is well approximated in 2D, and shape is clearly the main component strongly affected by TTD, shape will be the main focus of the Discussion.

### 4.1. Bringing 2D and 3D in the same data space: meaning, limitations and importance for shape

The second approach, that brings data into the same shape space, controlling for the bias between 2D and 3D, may not seem very intuitive. What is exactly the difference we have removed and why did we do it specifically for shape?

2D and 3D shape data, even if based on the same identical landmark configuration, belong to separate shape spaces of different dimensionality. Adding a zero Z coordinates to make a common superimposition possible is an expedient to bring them in the same Procrustes shape space. However, this is not enough to make the similarity relationships captured by each of type of data comparable. To appreciate why, one can look at Figure 8A, which shows a scatterplot of the first two PCs of shape of the combined data: 2D and 3D shapes are perfectly separated, and very distant one from the other, along PC1. The visualization of shape change along this axis, in correspondence of the mean of each cluster, shows that it is capturing the depth of the cranium, orthogonal to the ventral view (as it is particularly clear in side view, where the 2D shape is perfectly flat). For 2D data, this is missing information, as we cannot measure depth in a flat picture. This is clearly the major difference between the two types of data. Yet, what really matters for assessing TTD is whether, despite this lack of information, 3D similarity relationships are not distorted in 2D data. Crudely speaking and using an analogy, this is akin to having two scorers of an application for funds, with one who tends to be, for instance, consistently more positive than the other. Because this is a directional error, in order to make the assessment of the applicants comparable and check how well, despite it, the two sets of scores match, we might want to control for the bias. This can be achieved by subtracting the mean difference from the more positive set of scores or, which is equivalent, by mean centering the scores of both. Similarly, with the 2D/3D data brought in the same shape space by adding a zero Z coordinates to the 2D landmarks, the factor dominating the difference between them is the missing information in relation to the ‘thickness’ of the structure. As for the scorer example, mean centering controls for this bias without changing the relative differences within each dataset (the ranking of applicants by scorer one and two, or the similarity relationships of equids in 2D and 3D). Thus, with data mean centered in the same shape space (Fig. 8B), we can finally directly compare how well 2D corresponds to 3D.

**Fig. 8.**
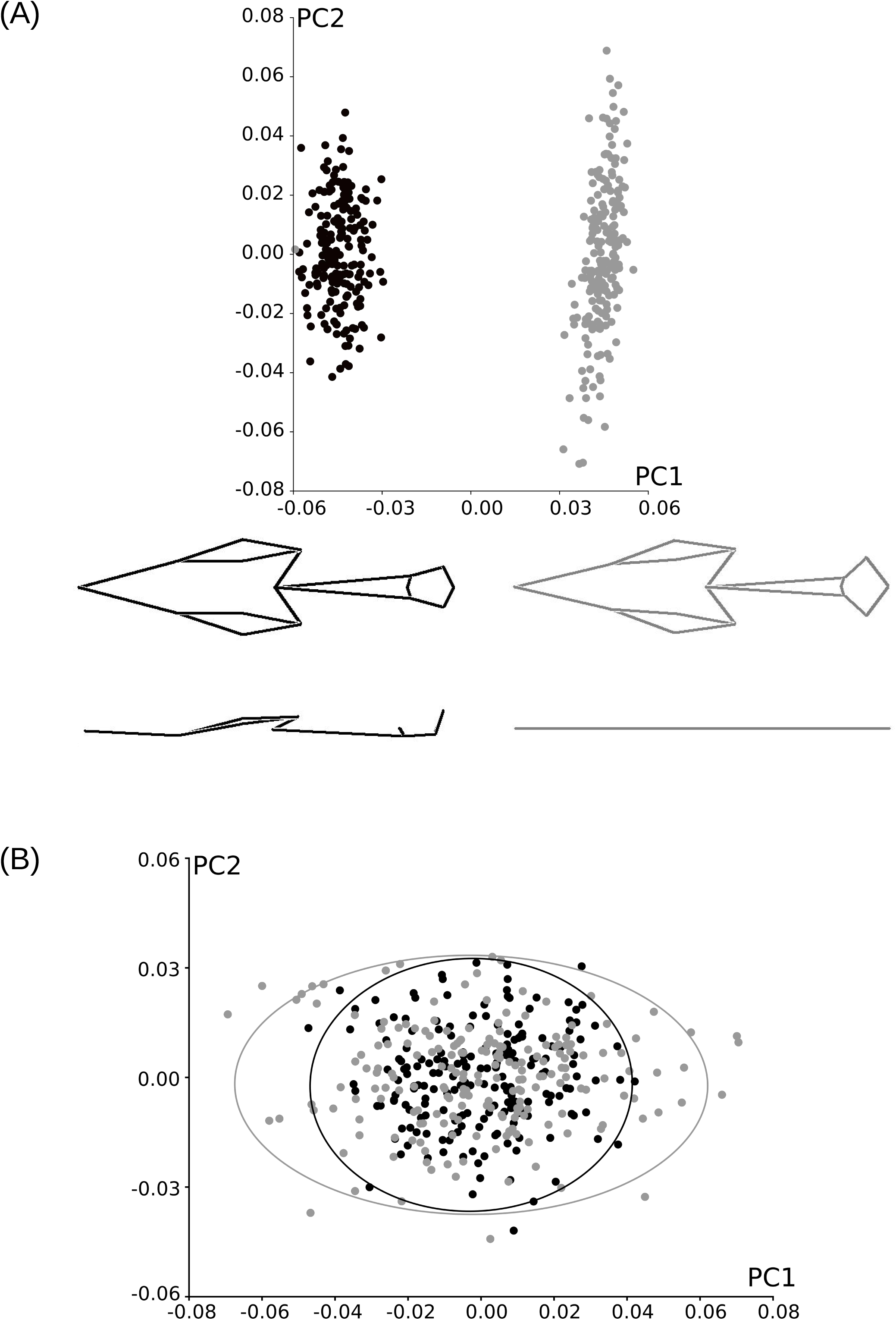
Common shape space analysis: scatterplots of PC1-PC2 of shape before (A) and after (B) mean-centering the 2D (grey colour) and 3D (black colour) data. Variance accounted for by PC1-2 are respectively 61.6% and 14.7% (A) and 37.6% and 15.0% (B). For (A), the ventral and side views of crania, corresponding to PC1 mean scores of 2D and 3D samples, are visualized below the scatterplot using wireframes (Klingenberg, 2013). In (B) 95% confidence ellipses are shown to emphasize the larger and more elongated pattern of variation in 2D compared to 3D.

This approach is simple, but has some limitations. A systematic difference is necessarily present between 2D and 3D data, but the method assumes that this bias is the same across all individuals and taxa. This may be true only as an approximation, and probably more likely when the depth (in the Z direction) of the different landmarks does not vary a lot across taxa. Also, the ‘common shape space’ approach may be effective to remove the bias, but it does not correct for differences in variance and covariance. These are another important aspect of morphological variation, that could be part of the assessment of TTD. We did not perform an specific test of similarities in variance and covariance (homoscedasticity), but the scatterplots of the first two PCs (Figs 3-4) seem to suggest that 2D shapes vary more than 3D ones.

A lack of homoscedasticity (i.e., heteroscedasticity) of 2D and 3D data may have an impact on tests relying on this assumption, such as the ANOVA in the common shape space, that should, therefore, be interpreted with caution. In subtle ways, in fact, differences in variance and covariance may affect also some of the analyses that do not assume homoscedasticity, such as the cluster analysis. The phenograms using individuals, unlike most other analyses, did suggest a rather poor TTD, and this could have been in part a consequence of heteroscedasticity. To understand why, one has to consider what happens if, compared to 3D, 2D shapes truly have more variance and a stronger covariance, as implicated by their larger and more elongated scatters on PC1-2 (Figs 3, 4 and 8B). If this really indicates heteroscedasticity, even if in relative terms inter-individual positions had been almost the same (e.g., specimen 1 relatively closer to 2 than to 3 in both datasets, and similarly for other individuals), 2D and 3D replicas of the same specimen could have ended up very distant one from the other in the shape space, and, therefore, in the phenogram. This is simply because, with heteroscedasticity in shape, the 2D space may be bigger or have, for instance, a longer PC1 and thus a different geometry. This type of variance-related problems is, using the simplistic analogy of the scorers, like having the same relative rankings but a larger spread of the scores by one of the two scorers. In our study, 2D shape variance is probably truly larger and thus a likely contributor to the differences between 2D and 3D shape relationships: unlike size, where variances were virtually identical in the two sets of data (not shown), total 2D shape variance was 8-27% (respectively at micro- and macro-evolutionary levels) larger than in 3D data (2D *vs* 3D shape variance in units of Procrustes distance in the common shape space: plains zebras, 0.00105 *vs* 0.00097; all species, 0.00137 *vs* 0.00118; species mean shapes: 0.00065 *vs* 0.00051).

Then, why is 2D shape variance in equid ventral crania larger than in 3D? We can only speculate about the reasons. For instance, it might be that 2D variance was inflated by the small but non-negligible inaccuracies in the standardization of the photos, because it is often hard to keep the same precise orientation of the camera relative to a study structure (Bakkes, 2017). If this is true, we would expect it to happen for most of the 2D data unless this aspect is extremely well standardized. This seems congruent with Cardini (2014), who found shape variance in 2D hemi-mandibles to be slightly larger than in 3D. However, this happened only for 2D data obtained using a flat-bed scanner, on which hemi-mandibles are difficult to be consistently positioned in the same way. In contrast, all other 2D data in that study used photos taken with a fairly sophisticated procedure (Cardini and Tongiorgi, 2003) to keep precisely the same orientation and distance of the camera to the structure. Indeed, these photos showed no increase in 2D shape variance, that was in fact even smaller than in 3D. If positioning errors are large and difficult to control for in the protocol for data acquisition, they could be reduced by a specific *a posteriori* mathematical manipulation of the landmark data (e.g., excluding the uniform component (Cardini, 2013) of shape?), but any correction of this kind would have to be carefully validated. In any case, in our study, heteroscedasticity may have contributed to the poor TTD in the phenograms of the individual specimens, but it is not its main explanation, as it did not happen with mean shapes (despite their largest relative difference in 2D/3D shape variance) and topologies in those trees differed even if computed separately for 2D and 3D (not shown).

Finally, in this first section of the Discussion, focusing on the ‘common shape space’ approach (Cardini, 2014), we would like to clarify better why it was only applied to shape. Part of the reason is that, with shape, there is no alternative in order to combine the 2D and 3D data and analyse them together in an ANOVA or a cluster analysis. However, if one is mainly interested in relative differences and there is a strong 2D to 3D bias also in size, this can be controlled for by mean centering, as for shape. In fact, with an accurate scale factor, we do expect 2D estimates of CS to be consistently smaller than 3D ones, because depth makes distances between 3D landmarks and their centroid larger. In our dataset, we did find this type of bias, but it was tiny: size was on average 10 mm larger in 3D (i.e., <2% of the mean 3D CS of 578 mm), and the variance explained by this difference was only 1.4%. Thus, we decided not to control for the TTD bias in size, whereas for shape it was crucial, as the bias accounted for 61% of variance in the common shape space. The reason why, unlike shape, CS showed a negligible 2D to 3D bias is probably simply that we carefully selected almost coplanar landmarks. The landmarks are, of course, not exactly on the same plane, but the differences in depth span at most a few centimeters, which is a very short distance in crania with an average condylobasal length of almost half a meter. Thus, the bias was negligible and TTD excellent for size.

### 4.2. Exploring the 2D approximation using the truss method

We already said at the beginning of the paper that, instead of trying to generalize from the few available studies, the best way to decide whether 2D data are appropriate for one’s own research is to preliminarily explore TTD in a subsample representative of the variation expected in the study. However, to do it, one needs 3D landmarks, that require either *ad-hoc* devices, such as potentially expensive 3D digitizers, or time and expertise for building detailed 3D models and digitize the landmarks on their surface.

There are, in fact, a few options to obtain 3D landmarks cheaply and relatively quickly. The R package stereomorph (Olsen and Westneat, 2015) allows to reconstruct 3D landmarks and outlines using two digital cameras and a calibration procedure to derive the 3D coordinates. After the calibration, the method should be efficient but requires not to change the zoom and position of the cameras, which might make it less suitable for specimens with a wide range of sizes. However, there is an even simpler low-tech/low-cost alternative to obtain 3D landmarks efficiently at least for small configurations of points. This is the reconstruction of 3D coordinates using linear inter-landmark distances and the truss method (Carpenter et al., 1996), available in Morpheus et al. (Slice, 1999), but also implementable in R (Claude, 2008). In Morpheus et al. the procedure is especially simple, as there is an option (“truss import”, in the file menu) to import linear distances, as well as a list of pairs of landmarks between which the distances were taken (both as simple txt files with renamed extensions, as explained in the “contents-trusses” of the help and exemplified in the TRUSS folder). To aid the 3D reconstruction, one can also add an optional template file with example landmark coordinates for a single individual, that can be made up by the user and will be later discarded. The approach has, nevertheless, some limitations and one may have to fiddle with the options and/or exclude a few individuals, if the algorithm fails to converge in the reconstruction. More importantly, the truss method becomes impractical for configurations with a large number of landmarks. However, to reduce measurements and save time, morphometricians can approximate the results from the full configuration by selecting a convenient subset of landmarks spanning the most important anatomical regions under investigation.

To provide an empirical example of this approach, we quickly explored its application in a sample of 30 randomly selected plains zebras, using five landmarks from the complete configuration of Figure 1 (i.e., landmarks 1, 5, 6, 10, 14). These landmarks capture the relative proportions of the main ventral cranial regions, which are the palate and cranial base. To apply the truss method, one should take caliper measurements between the 10 pairs of landmarks defined by those five points in the 30 individuals (in fact, with a careful selection, one could make the reconstruction using less inter-landmark measurements (Claude, 2008)). As this is an example, and the real specimens were no longer available to us for the caliper measurements, we simply computed the linear distances from the 3D Cartesian coordinates we already had and imported them in Morpheus et al. (Slice, 1999) as if they were actual caliper measurements. Finally, with the truss coordinates reconstructed and the corresponding 2D landmark data, we repeated some of the TTD tests in order to see if the results of the main study could be replicated with the ‘truss approach’ in a reduced dataset.

In the ‘truss dataset’, we found totally negligible sex differences, almost identical in magnitude (R^2^<1-2% for respectively size and shape) to those of the complete dataset (Table 2A). In both sexes allometry accounted for 11% of variance in 2D and 9% in 3D, which is about the same as in the complete dataset for females, but about twice bigger than that for males (Table 3A). Despite small differences, therefore, the ‘truss approach’ in a reduced dataset led to the same observation of very good congruence between 2D and 3D in relation to both sexual dimorphism and allometry. Correlational analyses also replicated well the findings of the main study (Table 4), with correlations between 2D and 3D estimates of size and shape of respectively 1.00 and 0.27. These are very similar to those from the main dataset for size, and only slightly lower for shape. Finally, individual variation in the ANOVA was, compared to differences between 2D and 3D data, two orders of magnitude bigger for size (R^2^s respectively of 99.5% and 0.1%) and three times larger for shape (R^2^s respectively of 75% and 24%), with the remaining tiny amounts of variance accounted for by sex. As in the previous analyses of the ‘truss dataset’, the ANOVA confirmed the results of the main study and, in fact, for shape produced almost identical estimates of R^2^s.

In summary, with few minor differences, we reached the same conclusions, as in the complete dataset, by using the ‘truss approach’ in a reduced set of data. This method, thus, seems a promising low-cost and time-saving expedient to explore the appropriateness of 2D data. As in all preliminary investigations using smaller datasets, a morphometrician will have to interpret its findings with caution. However, even a preliminary and simplified assessment of TTD will be better than a complete lack of it, which has been unfortunately the rule in 2D GMM until now.

### 4.3. Sampling issues? Heterogeneous N, smaller samples and number of landmarks

Comparative studies should ideally be done on samples of equal (or almost equal) size. That has a number of advantages in terms of design and statistics (for instance, it avoids issues with the choice of sum of squares in ANOVAs or with unbalanced samples in DAs (Kovarovic et al., 2011)). However, in zoology and especially with large animals from museum collections, one has to rely on what is available and try to maximize the output of data collection, which often depends crucially on the availability of funds (or, increasingly often in taxonomy, on a lack of them). The equids analysed in this paper are a good example of this problem: samples are very heterogeneous in size and certainly too small in some species to produce robust results (Cardini et al., 2015). This is less of an issue in a methodological study, whose main interest is to explore how well 2D data approximate 3D ones, but, even in this case, a potential effect of sampling on the accuracy and robustness of TTD results should be considered.

For instance, the large sample of plains zebras could have strongly influenced the outcome of the macroevolutionary analyses, as it accounts for about half of total N and is on average more than six times larger than those of other species. This is a genuine concern, but also one that, unlike problems with the small N of some other species, can be easily explored by repeating the analyses after selecting a random subsample of plains zebras. We did this (results not shown) using 16 specimens, which is the average sample size in the other species, and found a few minor differences, such as slightly larger correlations of 2D and 3D shape distances, as well as slightly inflated R^2^s (as expected in small samples (Cramer, 1987)). However, overall results were very similar to those of the total sample, suggesting robustness to this aspect of the highly unbalanced sampling of our study.

Total sample size may also be important in the assessment of TTD. Small samples produce unreliable estimates of means, variances and covariances, and shape data seem particularly sensitive to the problem (Cardini and Elton, 2007; Cardini et al., 2015). Small samples may also bias some of the results and, if this is less of an issue for R2, as it is inflated in both 2D and 3D data and therefore does not change the conclusion on TTD, an over- or under-estimate of matrix correlations, that also showed small differences in the analyses using subsamples of plains zebras, would be a concern. Indeed, in the small sensitivity analysis of the previous paragraph, using a random subsample of 16 plains zebras, matrix correlations were consistently larger than in the main study; yet, they were smaller in the subsample of 30 individuals used for the ‘truss approach’. This variability does not seem to suggest a clear bias, but it is only two examples and the ‘truss dataset’ may not be strictly comparable, because it included only ca. 1/3 of all landmarks. The distribution and number of landmarks is indeed another important aspect of TTD, but one that we did not investigate, because our configuration consists of too few landmarks. However, despite the obvious limitations, we briefly explored the sensitivity of 2D-3D shape matrix correlations to both sample size and the number of landmarks. As for the ‘truss approach’, this was not a main aim of our study, which is why the sensitivity analysis is concisely presented and discussed here.

We first focused on sample size. Thus, we extracted random subsamples of 5, 10, 25 and 50 individuals either in plains zebras or in the total sample with all seven species, and calculated matrix correlations between 2D and 3D shape distances. For each subsample, we repeated the analysis 500 times. The analyses showed that, although smaller samples had a larger range of correlations, both the mean and median of the 500 runs were virtually constant regardless of N (in plains zebras, mean r ranged from 0.43 in smaller subsamples to 0.46 in larger ones, whereas median r was 0.45-0.46 in all instances; in the total sample with all species, the mean ranged between 0.55 with N=5 and 0.61 with N=50, with the median being about 0.61 regardless of N). Thus, smaller samples do not seem to produce biased estimated of TTD, at least in terms of correlations between 2D and 3D shape data. However, as expected from De Moivre’s equation (Wainer, 2007), uncertainties increase in smaller samples and therefore the precision of TTD assessments is reduced.

After looking at the effect of sampling, we investigated the consequences of using fewer landmarks. We did this by selecting random configurations of six and nine landmarks from the total configuration, using a randomization that preserved pairs of bilateral landmarks. For each random configuration, we Procrustes superimposed the data and recomputed the correlations between matrices of 2D and 3D shape distances, both using only plains zebras or including the total sample of all 200 equids. For each sub-configuration, we replicated the random selection 500 times, which is enough to exhaust all possible combinations of six points and probably almost all those with nine landmarks. We found that, as with sample size, on average, having less landmarks did not seem to make a large difference: both mean and median correlations, using either six or nine landmarks, were about 0.4 and 0.55 respectively at micro- and macro-evolutionary levels. Nevertheless, compared with the slightly larger correlations from the main analysis including all 16 landmarks (respectively 0.47 and 0.61), it seems that having fewer landmarks could lead to slightly worse 2D approximations of 3D shapes. These results are, however, very preliminary: they may be specific to the dataset, and certainly are not robust enough to make generalizations. We clearly do not advise to increase the number of landmarks as a ‘quick fix’ for TTD problems. In fact, the choice of the landmark configuration must be hypothesis driven (Klingenberg, 2008; Oxnard and O’Higgins, 2011) and, in some analyses, more is not better (Cardini et al., 2019). The problem of a potential effect of the number and distribution of landmarks on TTD will have to be explored more extensively in future studies using larger sets of points and ideally many taxonomic groups and structures.

### 4.4. Conclusions: 2D or not 2D?

The question from the title of another TTD study (Buser et al., 2018) (p. 806), “2D or not 2D?”, looks like a clear dichotomy, but the answer is not a simple “yes” or “no”. The picture emerging from TTD research is complex. In our study, for instance, it is evident that 2D data approximate 3D size very well, but are less good at quantifying shape, despite the selection of almost coplanar landmarks. However, there are several nuances and some apparent contradictions.

As in Cardini (2014), 2D cranial data appear to be inaccurate at capturing the fine details of shape relationships at micro-evolutionary level in homogeneous samples (same age class). In fact, although the range of correlations between 2D and 3D shape distance matrices is similar in these studies (and also similar to those of Cardini and Thorington’s (2006) 2D and 3D ontogenetic analysis of marmots), the percentage of 2D shapes clustering with their 3D replica in the phenograms is much lower in the equids (<10%) than in marmots (37-85% – Cardini 2014). The equid sample is in fact four or more times larger than any of the marmot samples, and this is something that may have increased the chances of a mismatch between 2D and 3D shapes. Yet, even in marmots, despite their smaller samples and with the exception of the fairly flat hemi-mandibles, the 2D-3D mismatch in the phenograms occurred in about half of the individuals, which clearly does not suggest a good TTD. Overall, these are worrying results, because intraspecific variation is a main field of study in morphometrics.

With larger differences, however, such as those among different species, TTD improves and may be thus less of a concern at supraspecific level, as well as in many ontogenetic analyses of large age-related morphological differences (Cardini and Thorington, 2006). Can we, therefore, conclude that TTD is problematic only when measuring small amounts of biological variation? Some of the research summarized in the Introduction suggests potential problems even with macro-evolutionary analyses (e.g., (Buser et al., 2018; Hedrick et al., 2019)). In our work too, although TTD improves at supra-specific level, it is still large especially if judged in terms of the not so high correlations between shape matrices (ca. 0.6-0.7) and the large R^2^ of measurement error (ca. 1/5 of total shape variance). Besides, the percentage of 2D-3D replicas of an individual clustering as ‘sisters’ in the phenogram of all 200 equids is no better than using only plains zebras. This again suggests poor accuracy in the description of subtle inter-individual relationships and seems to lead to the conclusion that 2D ventral cranial shape data in equids are in fact problematic at all levels, micro- as well as macro-evolutionary.

However, TTD must be assessed in relation to the hypotheses under investigation. In this respect, despite the substantial differences in details, answers to the main biological questions we used as examples in our study are almost identical in 2D and 3D: sexual dimorphism in plains zebras is negligible (R^2^<2%); species differences are present and large (accounting for ca. ¼ to more than ¾ of variance in respectively shape and size); and patterns of group and allometric variation are very similar (Figs 2-5 and 7). Moreover, a strong congruence between the 2D and 3D results is also supported by the angles between 2D and 3D main vectors of variation in the common shape space, which, for the large majority, are small. For instance, for the first two PCs, they are 16° or less in the total sample, 19-32° using mean shapes and 34-59° including only plains zebras; for the allometries, angles between 2D and 3D vectors are 27-34° in respectively females and males of plains zebras and 25° using species means.

Thus, unlike for instance (Hedrick et al., 2019), who reported important differences in 2D *versus* 3D results (e.g., in asymmetric changes, as well as group differences, found only in 3D), we reached almost identical conclusions on patterns of ventral cranial variation in equids using either 2D or 3D analyses. This seems paradoxical given the clearly non-negligible differences in the fine details of inter-individual shape variation (the modest matrix correlations, large R^2^s of 2D-3D differences, and the inaccuracies in phenograms, that we mentioned above). However, similarities in main patterns despite variability in precise inter-individual differences can be reconciled, if the distortion of 2D shapes is mostly occurring in non-patterned components of shape variance. This would mean that the 2D approximation of ventral crania in living equids increases the measurement error in the data, but does so in a random fashion, which makes the data ‘noisier’ without significantly perturbing the main signal of morphological variation in relation to sex, species, and allometry.

As most of the time morphometricians focus on broad patterns, our finding of very good congruence in the results of the main biological hypotheses we tested provides a degree of optimism on the possibility of an effective use of 2D data to approximate variation in 3D biological structures. As pointed out by Navarro and Maga, 2016, although 3D shape is more accurate, 2D data might often correctly capture the main big picture of evolutionary and developmental change. Nevertheless, for now, we must also acknowledge that there are too few studies to draw a solid conclusion and, therefore, the question of the appropriateness of 2D analyses of 3D shapes is still open.

To further stress that any generalization is premature and cannot be based on a small amount of case-specific evidence, we end the Discussion by comparing the only two studies we know, Álvarez and Perez (2013) and Cardini (2014), that analysed TTD using the same structure (hemi-mandibles) in the same order of animals (the rodents). Álvarez and Perez (2013) investigated shape variation across different genera of caviomorph rodents and found general agreement in 2D and 3D macro-evolutionary patterns, but also some differences, with dissimilarities especially pronounced when they restricted the analyses to intrageneric variation. Cardini (2014), in contrast, using a fairly similar set of landmarks on marmot hemi-mandibles, suggested that the 2D flattening may be much less problematic in supraspecific analyses. The two studies may not be strictly comparable because of differences in methods, as well as some of the landmarks. Yet, if TTD errors in interspecific comparisons of caviomorph hemi-mandibles are truly larger than in marmots, this may largely depend on the lateral flaring of the hystricognath hemi-mandible, that makes it more 3D than those of sciurognaths. This again suggests that the accuracy of 2D data may be influenced by a complex range of factors, that do not only include the magnitude of the differences, but also landmark coplanarity and taxon-specific anatomical variability.

Openshaw et al. (2017) suggested that “studies restricted to a 2D geometric morphometric analysis of a complex 3D biological structure can combine carefully designed 2D landmark configurations describing alternative planes [e.g., ventral, side and dorsal views] to maximize shape coverage”. That may be true, but could also, as in many other cases, be specific to their study. Using different 2D views may indeed increase the amount of information in the data, but does not guarantee that any of those views is a good approximation of the 3D anatomy. More generally, morphometricians should rely less on untested assumptions and more on objective assessments of measurement error (including TTD) in their own data and in relation to the specific study hypotheses. This means that one should seriously consider preliminary exploratory analyses, before investing energy, money and time in extensive collections of 2D data, whose accuracy may or may not be adequate for the specific aim, structure and taxon.

## Acknowledgements

Data were collected at Museo Civico di Storia Naturale di Milano, Natural History Museum Vienna, National Museum Prague, Museum für Naturkunde, Museum National d’Histoire Naturelle, Hungarian Natural History Museum, Naturhistoriskariksmuseet. We owe a huge thank to all curators, collection managers and staff, including the SYNTHESYS administrators, whose help and support was fundamental. Among them, a special thanks to: Giorgio Bardelli, Görföl Tamás, Gabor CSORBA, Virginie Bouetel, Aurelie Verguin, Frank Zachos, Alex Bibl, Krapf Andrea, Petr Benda, Karel Kaderàbek, František Vacek, Manja Voss, Christiane Funk, Detlef Willborn, Mayer Frieder, Daniela Kaltoff, Irene Bisang, and Emily Dock Åkerman. A special thank also to the greatly missed Colin Groves, for his fundamental contribution to interpreting the preliminary analyses of wild asses.

## Author contributions

The study was designed and the paper written by AC; 3D data were acquired and pictures were taken by AC; 2D landmarks were digitized by MC with AC supervision; data were analysed mainly by AC with preliminary analyses by MC. *Competing interests:* We declare no competing interests or conflict of interest.

